# Direct cryopreservation of poultry/avian embryonic reproductive cells: A low-tech, cost-effective and efficient method for safeguarding genetic diversity

**DOI:** 10.1101/2021.09.28.462137

**Authors:** Tuanjun Hu, Lorna Taylor, Adrian Sherman, Christian K. Tiambo, Stephen J. Kemp, Bruce W. Whitelaw, Rachel J. Hawken, Appolinaire Djikeng, Mike J. McGrew

## Abstract

Chickens are an important resource for smallholder farmers who raise locally adapted, genetically distinct breeds for eggs and meat. The development of efficient reproductive technologies to conserve and regenerate chicken breeds safeguards existing biodiversity and secures poultry genetic resources for climate resilience, biosecurity, and future food production. The majority of the over 1600 breeds of chicken are raised in low and lower to middle income countries (LMICs) under resource limited, small scale production systems, which necessitates a low tech, cost effective means of conserving diversity is needed. Here, we validate a simple biobanking technique using cryopreserved embryonic chicken gonads. The gonads are quickly isolated, visually sexed, pooled by sex, and cryopreserved. Subsequently, the stored material is thawed and dissociated before injection into sterile host chicken embryos. By using pooled GFP and RFP-labelled donor gonadal cells and Sire Dam Surrogate (SDS) mating, we demonstrate that chicks deriving entirely from male and female donor germ cells are hatched. This technology will enable ongoing efforts to conserve chicken genetic diversity for both commercial and small holder farmers, and to preserve existing genetic resources at poultry research facilities.

## Introduction

Chickens, with a global population over 60 billion, are the most populous bird species on the planet (Ritchie and Roser 2021). Regionally adapted chickens (considered as indigenous breeds or local ecotypes) are found in every country and are genetically diverse and well adapted to scavenging feeding, environmental challenges and climatic conditions (DAD-IS 2021). As rural farming practices become replaced by centralised commercial poultry breeding, local chicken breeds are at risk of becoming extinct. This loss of local ecotypes with their unique genetic diversity jeopardises future improvements in livestock climate adaptation and sustainable farming practices (Alders and Pym 2009, Melesse 2014). A conservation programme integrating both DNA sequencing and reproductive biobanking of local chicken breeds and ecotypes would provide a platform for protecting and conserving the genetic diversity of chicken and also serve as an exemplar for other domestic poultry species (Woelders, Windig et al. 2012, Whyte J. 2015). Chickens are also a model system to study development, avian immunology and diseases (Davey, Balic et al. 2018). The hundreds of research chicken lines kept at avian facilities are also at risk of loss (Fulton and Delany 2003).

The reproductive cells of an animal, the germ cells, contain the genetic information that is transferred from one generation to subsequent ones. The differentiated germ cells, the highly specialised sperm and egg, each carry a haploid genome and recombine to form the diploid fertilised egg. Cryopreservation of adult germ cells in ruminant livestock is now routine, however, in avian species, cryopreservation of the mature gametes is problematic. The large yolk-filled bird egg cannot be cryopreserved. Cryopreserved chicken semen has poor fertilisation rates for some chicken breeds when used in artificial insemination (Thelie, Bailliard et al. 2019). This may be due in part to the length of the avian oviduct and the prolonged storage of semen in specialised glands of the oviduct, the deleterious effects of the cryopreservation and thawing process, and the contraceptive effects of the cryoprotectants (e.g. glycerol) in the freezing media. (Blesbois, Grasseau et al. 2008, Matsuzaki, Hirohashi et al. 2021).

Avian embryonic germ cells, in contrast, can be efficiently cryopreserved. The avian germ cell lineage originally consists of ∼40 diploid cells in the laid egg (Karagenc, Cinnamon et al. 1996). These cells, the primordial germ cells (PGCs), migrate through the vascular system of the developing embryo and colonise the forming gonads. The gonadal germ cells develop into the terminally differentiated oocytes numbering over 100,000 meiotic follicles in the hatched female chick and the proliferative spermatogonial stem cell population of the male testis. We, and others, have demonstrated that PGCs can be isolated from individual embryos and propagated *in vitro* in a defined cell culture medium to produce several 100,000 cells in 3-4 weeks (van de Lavoir, Mather-Love et al. 2006, Whyte, Glover et al. 2015). This population of PGCs can then be cryopreserved in multiple aliquots. The cryopreserved PGCs are later injected into the vascular system of surrogate host embryos (∼3000 PGCs/embryo). The gonads of the host embryos are colonised by the exogenous germ cells and, when mature, will produce functional donor PGC-derived gametes. Chemical or genetic ablation of the host embryo’s endogenous germ cells will increase the frequency of offspring formed by the exogenous germ cells (Nakamura, Yamamoto et al. 2008, Macdonald, Glover et al. 2010). When hens genetically modified to disrupt the germ cell determinant, *DDX4*, were injected with donor cryopreserved female PGCs from a different breed of chicken, they laid eggs that were solely derived from donor cells (Woodcock, Gheyas et al. 2019). Similarly, transgenic chickens expressing an inducible Caspase9 transgene in the germ cell lineage were treated with the dimerization chemical, AP20187 (B/B), to ablate the endogenous germ cells. iCaspase9 surrogate hosts of both sexes produced gametes and offspring only deriving from donor PGCs. Direct mating of the surrogate hosts (Sire Dam Surrogate (SDS) mating) produced genetically pure breed offspring entirely derived from the exogenous PGCs ((Ballantyne, Woodcock et al. 2021), Ballantyne, in press).

The *in vitro* propagation of PGCs is technically demanding, expensive, and requires a complex cell culture medium and specialised cell culture facilities. The *in vitro* propagation of PGCs leads to epigenetic modifications and reduced germ line transmission with the increased period of *in vitro* culture ((Woodcock, Gheyas et al. 2019); Ballantyne et al, in press, Govoroun et al, in press). Thus, a biobanking methodology that did not depend on culture of germ cells would be preferable for poultry conservation. To address these current limitations, we investigated the use of directly cryopreserved embryonic gonadal germ cells for the cryoconservation of chicken breeds. The initial PGC population rapidly increases after colonising the gonadal anlagen and numbers many tens of thousands of mitotically active cells by the middle of the incubation period. This proliferative phase is followed by the mitotic quiescence of male germ cells until hatch and the meiotic entry of female gonadal germ cells at embryonic day (ED) 15 (Hughes 1963). Male and female gonadal germ cells re-introduced into migratory stage embryos will re-migrate and colonise the host gonad (Tajima, Naito et al. 1998, Naito, Minematsu et al. 2007). In fact, researchers have previously demonstrated that gonadal germ cells will colonise a host gonad and form both functional gametes and offspring (Tajima, Naito et al. 1998, Mozdziak, Wysocki et al. 2006). The gonadal germ cells should therefore be adequate for biobanking and re-establishing a chicken breed if combined with a sterile host bird.

Here, using fluorescently labelled reporter lines of chicken, we confirm that the chicken gonad between ED 9 and 12 contains over 10,000 germ cells and that gonadal germ cells up to ED 10 of incubation are capable of efficient re-migration to the gonad when injected into the vascular system of an ED 2.5 host embryo. The ED 9 chicken gonad (stage 35 HH) can be visually sexed, pooled, and directly cryopreserved using a simple freezing medium before long-term storage in liquid nitrogen (Fig. 1A). Subsequently, the cryopreserved gonads are thawed and dissociated before introduction into iCaspase9 host embryos. Male gonadal cells are directly injected into male sterile surrogate host embryos. Female gonadal cells are MACS purified to enrich for the germ cell population then injected into female surrogate host embryos. Using GFP- and RFP-labelled gonadal cells, we show that multiple donor genotypes are transmitted through the male and female hosts and ‘homozygous’ donor-derived offspring can be directly generated through SDS mating (Fig. 1B). Application of this technology and future variations should enable economical and efficient biobanking of indigenous poultry breeds in LIMCs and be an exemplar for avian species cryo-conservation.

**Fig. 1.**
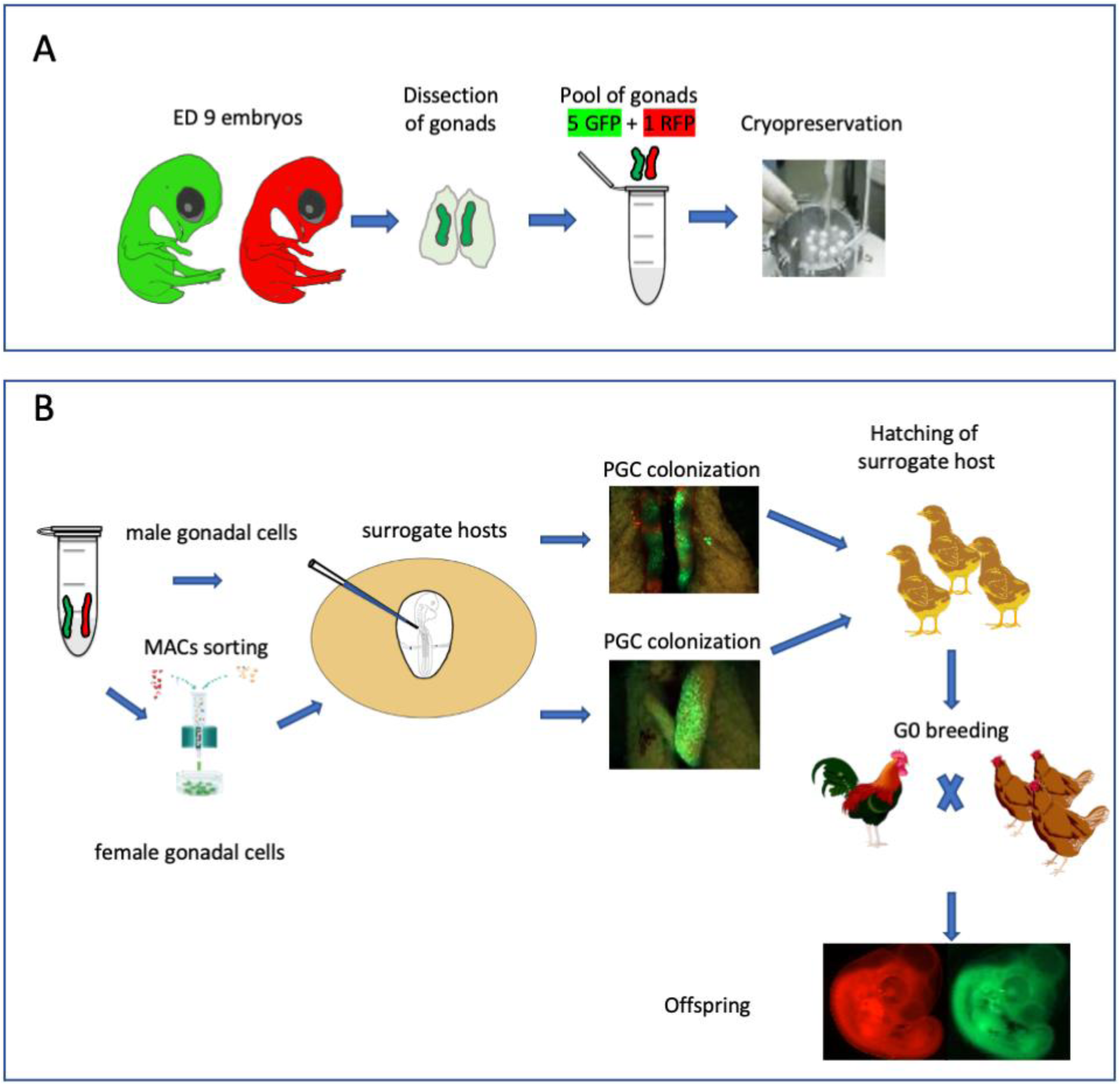
Isolation and cryopreservation of embryonic gonads followed by transmission through sterile surrogate hosts A. ED 9 gonads are isolated from embryos, pooled by sex, and cryopreserved in liquid N_2_. B. The frozen gonads are thawed, dissociated and injected into sterile surrogate host embryos. The surrogate host embryos are incubated and hatched and bred to hatch donor gonadal offspring.

## Results

### The germ cell population of the avian embryonic gonad

We previously generated an iCaspase9 chicken line that contained an inducible Caspase9 gene and a GFP reporter integrated in the chicken *DAZL* locus (Ballantyne, Woodcock et al. 2021). We demonstrated that this GFP reporter construct was expressed exclusively in all germ cells of the developing embryo. We used this iCaspase9 GFP reporter gene to quantitate the germ cell population in the chicken gonad from ED 9 to ED 12 of development (Fig. 2A). We chose ED 9 as the starting point as it is the earliest developmental stage that the sex of male and female gonads can be clearly distinguished by visual inspection. Between ED 9 and ED 12 total cell number in the female gonad increased 8-fold. The population of female germ cells in the gonad increased more than 40-fold during this period and the percentage of female germ cells rose from 1.9% to 10.6%. In contrast, in the male gonad the total gonad cell number increased 3.6-fold between ED 9 and ED 12 and the germ cell number rose 3.1-fold. The proportion of male germ cells remained between 2.4-2.9% during this period. These observations are similar to those previously reported using a different quantitation technique (Yang, Lee et al. 2018). The total gonadal germ cell population, therefore, is greater than 10,000 cells in both sexes by ED 10.

**Fig. 2.**
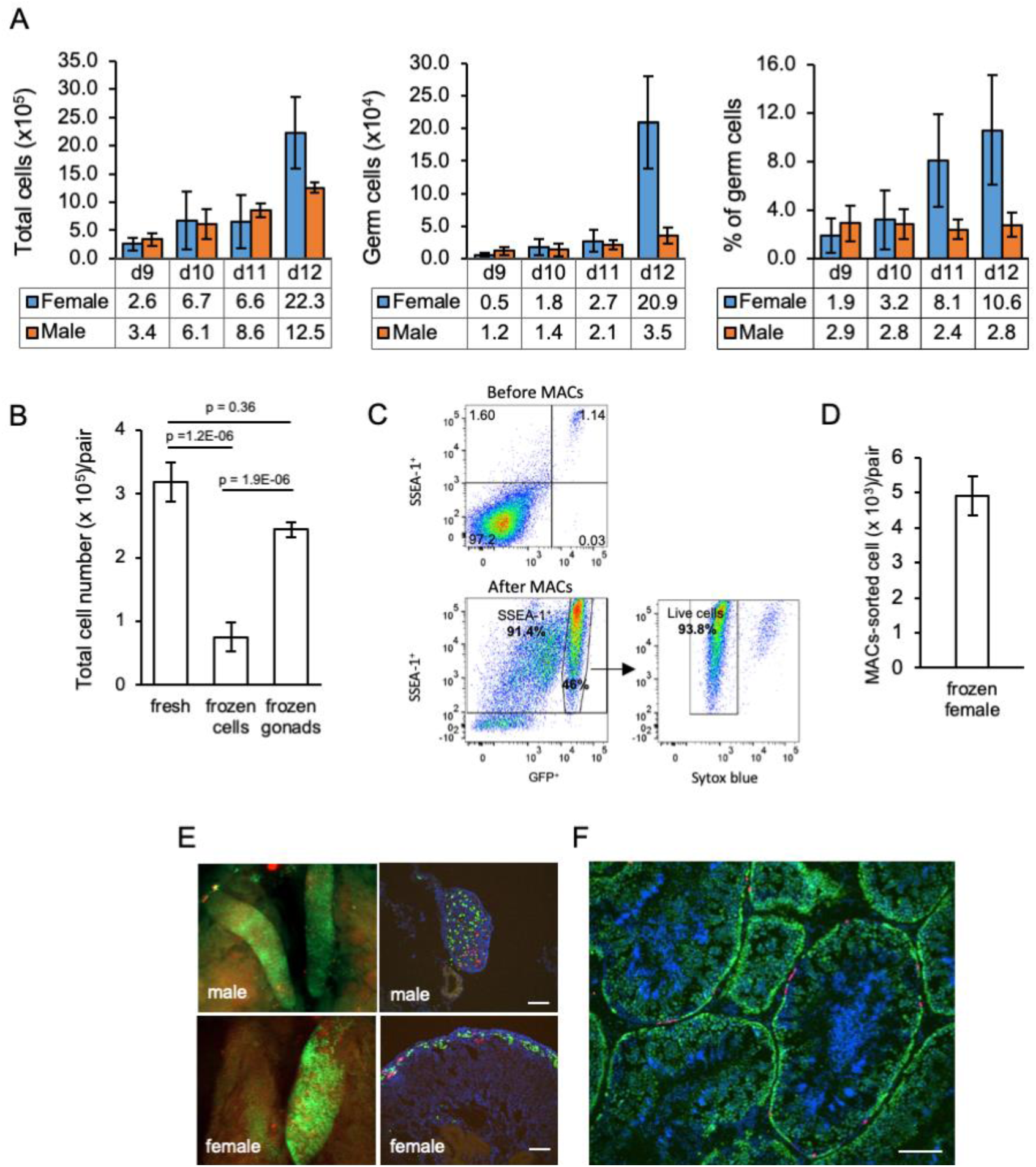
Characterisation and cryopreservation of gonadal germ cells A. Population of gonadal germ cells between ED 9-12. The number of germ cells were determined by their expression of GFP protein in iCaspase9 transgenic embryos (n = 3-7 gonad pairs for each sex at each day). B. Yield of viable dissociated cells directly from freshly isolated embryonic day 9 gonads (control), cryopreserved dissociated gonadal cells subsequently thawed (frozen cells), and cryopreserved whole gonads subsequently thawed then dissociated (frozen gonads). Cell viability was determined using a trypan blue exclusion assay. Data from 13-20 independent experiments using mixed male and female gonads. C. Flowcytometric analysis of MAC sorted female gonadal cells. Frozen female ED 9 (HH35) gonads from iCaspase9 GFP^+^ embryos were MAC sorted using an anti-SSEA-1 antibody. The purified cells were then immunostained by secondary antibody to SSEA-1 to detecting the percentage of SSEA-1 cells expressing GFP. The GFP^+^ population was analysed for viability using Syto blue; n = 5 independent experiments. D. Yield of MAC sorted cells from cryopreserved ED 9 (HH35) female gonads. The average number of GFP^+^ cells purified by MACS from a single iCaspase9 embryo using an anti-SSEA-1 antibody. Data from 5 independent experiments using 12-26 gonad pairs per experiment. E. Colonisation of sterile iCaspase9 embryos by cryopreserved male and female gonadal cells. The host ED 14 gonads are shown on the (left) and transverse sections from those gonads are on the (right). Day 2.5 iCaspase9 host embryos were injected with gonadal donor cells at a 5GFP^+^:1RFP^+^ ratio mixed with B/B compound; 45,000 male cells/embryo. Female cells were MAC-sorted before injecting 1,400 female cells/embryo. Representative embryo shown (n > 5, for each sex). Scale bar = 100um. F. Seminiferous tubule of an adult testis (>6 months) from a sterile surrogate host injected with donor male cells prepared as in C. Representative testes section from n = 7 males. Scale bar = 100um.

**Fig. 3.**
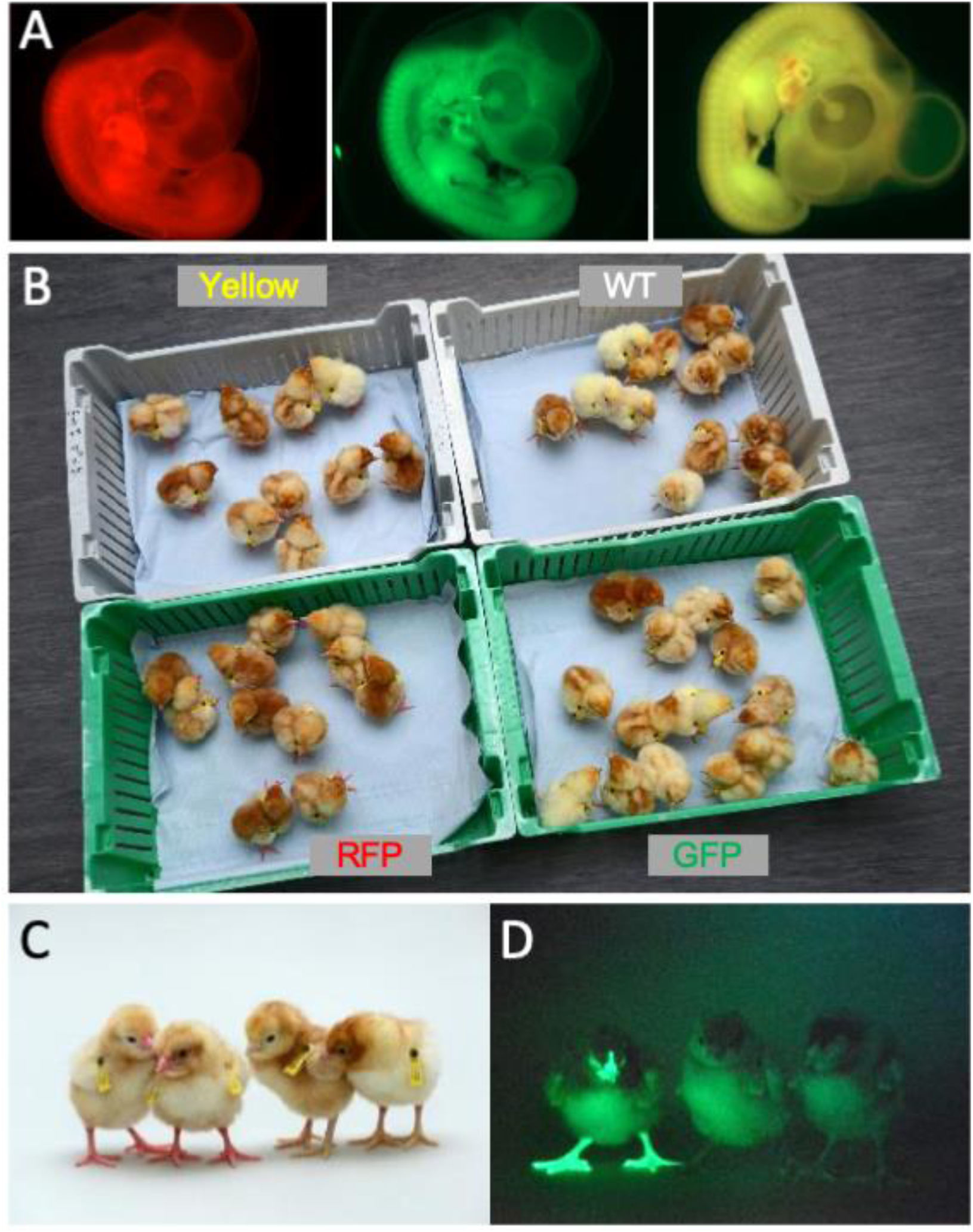
Hatchings from ♂ surrogate hosts injected with ♂ gonadal cells mated to ♀ surrogate hosts injected with ♀ gonadal germ cells A. ED 5 embryos SDS mating displaying representative red, green, and yellow florescence. B. Hatched chicks from SDS mating grouped according to fluorescence C. RFP florescent chicks were apparent (chicks on left) under visible light D. GFP fluorescent chick (left) visualised under GFP illumination

### Cryopreservation of embryonic gonads and recolonization of host embryos

We assayed for gonadal cell survival after cryopreservation and thawing. Gonadal cell viability was severely reduced after the cryopreservation of dissociated gonadal cells (Fig. 2B). In comparison, cryopreservation of whole gonads followed by thawing and subsequent cell dissociation did not significantly reduce gonadal cell viability when compared to the viability of directly dissociated cells from freshly isolated gonads (Fig. 2B). We next assayed the capacity of cryopreserved gonadal cells to re-colonise the gonad of surrogate host embryos using GFP^+^ and RFP^+^ labelled gonads from ED 8 to ED 11 transgenic embryos as donors. The migration of donor gonadal germ cells to a host gonad can be easily quantified by counting the fluorescent cells in the host gonads. Whole donor gonads were cryopreserved and subsequently dissociated cells prepared from these frozen gonads were injected into wildtype host embryos. We injected 10,000-15,000 female gonadal cells (∼150 germ cells) into the host embryos and observed germ cell colonization 5-6 days (ED 8-9) post injection. Gonadal cells from ED 8-10 donor female embryos achieved repeatable colonisation of host gonads, while injection of ED 11 donor resulted in much fewer fluorescent cells present in the host embryo (Supplementary Fig. 1).

Male gonadal germ cells form the spermatogonial stem cell (SSC) population of the testes that proliferates and differentiates into spermatozoa for the entire life of the cockerel. In contrast, female gonadal germ cells enter into meiosis starting at ED 15 reaching a final population of 480,000 post-replicative germ cells in the hatched chick (Hughes 1963). We postulated that the number of fluorescent cells observed colonising the host female gonad (Supplementary Fig. 1) would not be sufficient to form an appropriate oocyte hierarchy in the mature ovary and a continuous egg laying cycle lasting the 10-12 month period of a typical layer hen. The cell surface stem cell marker, SSEA-1, is highly expressed on the surface of migratory and post-migratory gonadal germ cells until ED 9-10 of incubation after which expression of SSEA-1 is rapidly lost from the germ cell (Urven, Erickson et al. 1988). Selective enrichment of gonadal germ cells using SSEA-1 antibody was shown to increase the colonisation of the host gonad (Kim, Kim et al. 2004, Mozdziak, Angerman-Stewart et al. 2005, Mozdziak, Wysocki et al. 2006). We used Magnetic Activated Cell Sorting (MACS) to purify SSEA-1-expressing cells from the ED 9-10 (stage 35 HH) female gonads. We first determined that SSEA-1 expression on female gonadal germ cells decreased after ED 10 (Supplementary Figure 2). Cryopreserved iCaspase9 gonadal tissues were thawed and dissociated and MACS using SSEA-1 antibody was used to enrich for female PGCs. We observed that the SSEA-1 antibody captured and enriched >40-fold GFP^+^ gonadal germ cells (1.17% > 46%). The MACS purified cells were 94% viable and as expected co-expressed SSEA-1 antigen on their surface (Fig. 2C). The yield of germ cells from the female ED 9 gonad by MACS was approximately 5000 putative germ cells per pair of gonads (Fig. 2D).

We next tested the colonisation of male dissociated gonadal cells and MACS-enriched female gonadal cells using iCaspase9 host embryos treated with B/B compound. We used GFP^+^RFP^+^ gonad pairs mixed in a ratio of 5:1 to identify multiple colonisation events. A cell suspension of male gonadal cells or MACS purified female gonadal cells was mixed with B/B dimerization chemical and injected into the vascular system of ED 2.5 (stage 16 HH) iCaspase9 embryos (Fig. 2E). Whole tissue imaging and cryosections of surrogate host ED 14 gonads indicated that the donor PGCs from frozen gonadal tissues colonized the iCaspase9 host embryos of the same sex. The majority of cells were GFP^+^ and fewer RFP^+^ PGCs were also present in the host gonads (Fig. 2E). An analysis of cryosections of adult male testes indicated that the majority of putative germ cells in the seminiferous tubules were GFP^+^ and a minority of cells were RFP^+^ (Fig. 2F).

### Generation of iCaspase9 surrogate host chicken

Based on these preliminary results, we proceeded with hatching of iCapase9 surrogate host chicken injected with donor cryopreserved gonadal cells in order to measure germ line transmission rates of the donor material. Male dissociated gonadal cells were mixed with B/B compound and injected directly into ED 2.5 iCaspase9 embryos. B/B compound activates the iCaspase9 transgene leading to the selective ablation of the endogenous germ cells (Ballantyne, Woodcock et al. 2021). Female embryonic gonads were thawed, dissociated and MACS-purified using an anti SSEA-1 antibody before injection. We mixed GFP^+^ and RFP^+^ donor gonadal material in a 5:1 or 5:2 ratio in order to identify the transmission of multiple donor genotypes from individual surrogate host chickens. In our first experiment (CRYO-1M) the survival of injected embryos was >60% and the hatchability was >40% (Table 1). In subsequent experiments, the incubation conditions were altered (see Materials and Methods) and hatchability increased to 58-83%. From these injection experiments, we estimate that cryopreserved gonads from 6 ED 10 embryos would provide sufficient cryopreserved material to inject 25 host embryos from which 10 host chicks (5 male and 5 female) could be hatched. Furthermore, the mating of iCaspase9 homozygous cockerels mated to wildtype hens produced 100% heterozygote iCaspase9 host eggs for injections (CRYO1-4). In contrast, mating a heterozygote DDX4 Z^+^Z^-^ heterozygote cockerel to wildtype hens (CRYO-5F) produce eggs of which 25% were of the correct Z^-^W host genotype.

**Table 1.**
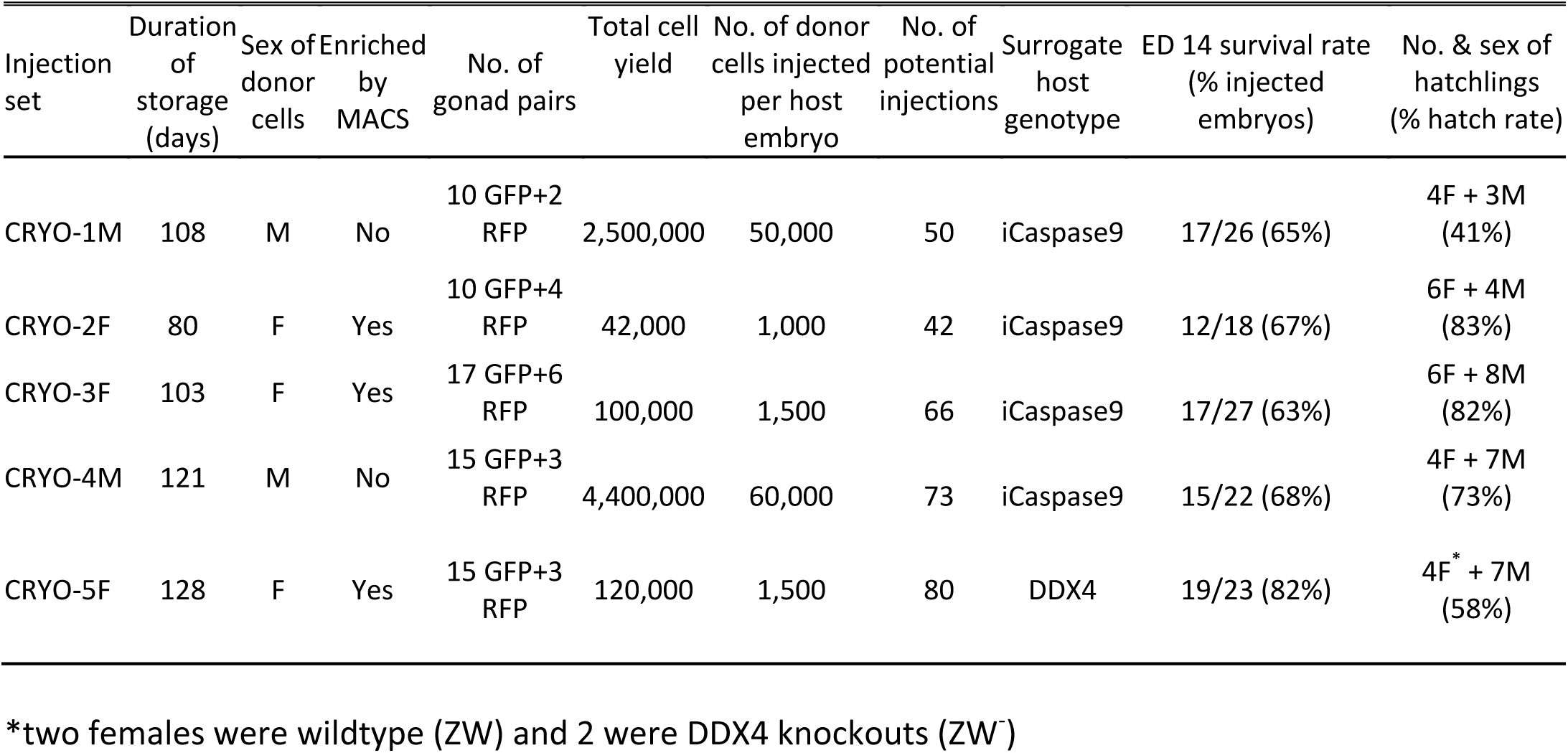
Injection and hatching of surrogate hosts

### Transmission of donor gonadal germ cells through sterile surrogate hosts

We established breeding groups of individual surrogate host cockerels mated to a cohort of wildtype hens and two cohorts of surrogate host hens mated to a wildtype male cockerel. We then assayed fertility of the breeding groups, the number of RFP^+^ and GFP^+^ offspring as a proxy for genotype transmission, and the presence of any iCaspase9 offspring which would indicate that the sterilisation of the surrogate host animal was not complete.

We generated two cohorts of iCaspase9 host males each injected with independently cryopreserved male gonadal samples (CRYO-1M, CRYO-4M) (Table 2). Individual iCaspase9 males were naturally mated to a cohort of wildtype hens and laid eggs were assayed for fertility and hatchability. Fertility was high for all males and ranged from 83-99%. We used the detection of the GFP and RFP reporter genes in the progeny as a proxy for measuring the transmission of multiple genotypes through the host males. The presence of both GFP^+^ and RFP^+^ embryos was observed for each male indicating that multiple donor gonadal germ cells formed functional spermatozoa in each male surrogate host. We assayed the embryos for the presence of the iCaspase9 transgene carried by the male surrogate host. 6 out of 7 host males did not transmit the endogenous transgene (0/643 embryos) suggesting that all embryos sired by these males derived from donor gonadal germ cells. 1 out of 7 host males transmitted the iCaspase9 transgene to 6% of the embryos (9/147) indicating that approximately 12% (the transgene was heterozygote in the surrogate hosts) of the embryos were derived from endogenous germ cells and 88% from exogenous donor gonadal germ cells. Analysis of the testes from the 7 cockerels showed normal germ cell development and differentiation (Supplementary Fig. 3A, 5A).

**Table 2.** Germline transmission from ♂ surrogate hosts injected with ♂ gonadal PGCs

| Host mating groups | No. of eggs laid per week * | Eggs set | Fertility<sup>§</sup> (% eggs set) | No. of RFP embryos (% fertile) | No. of GFP embryos (% fertile) | Transmission of surrogate iCaspase9 transgene (% fertile) |
| --- | --- | --- | --- | --- | --- | --- |
| CRYO-1M_1181♂ × 6 WT♀ | 6.7 | 147 | 96% | 4 (3%)<sup>&</sup> | 101 (72%) | 0 |
| CRYO-1M_1182♂ × 6 WT♀ | 6.6 | 148 | 99% | 9 (6%) | 89 (61%) | 9 (6%) |
| CRYO-1M_1179♂ × 6 WT♀ | 6.6 | 187 | 98% | 51 (28%) | 55 (30%) | 0 |
| total |  | 482 | 98% | 64 (14%) | 245 (52%) | 9 (2%) |
| CRYO-4M_1420♂ × 5 WT♀ | 4.5 | 89 | 91% | 4 (5%) | 64 (79%) | 0 |
| CRYO-4M_1424♂ × 5 WT♀ | 4.1 | 60 | 93% | 3 (5%) | 41 (73%) | 0 |
| CRYO-4M_1427♂ × 6 WT♀ | 6.9 | 74 | 85% | 13 (21%) | 29 (46%) | 0 |
| CRYO-4M_1430 ♂ × 6 WT♀ | 6.7 | 143 | 83% | 14 (12%) | 80 (68%) | 0 |
| Total |  | 366 | 87% | 34 (11%) | 214 (67%) | 0 |

<sup>§</sup>Fertility was assessed between ED 4-7
\*Lay rate; eggs were counted over a 60 day period when hens were between 7-12 months of age and divided by the number of fertile hens present in pen. The maximum possible lay rate is 7.0 eggs per hen per week.

We generated two cohorts of iCaspase9 surrogate hens each injected with independent cryopreserved gonadal samples (CRYO-3F, CRYO-4F) (Table 3). We also generated one cohort of DDX4 surrogate hens. The surrogate host hens laid between 5.0-6.3 eggs per week (out of a potential 7.0 eggs per week). This lay number was comparable to eggs laid by control wildtype brown layer hens: 4.1-6.9 eggs per week (Table 3). The female cohorts were naturally mated to wildtype cockerels and fertility of laid eggs ranged from 80-98%. The presence of both GFP^+^ and RFP^+^ embryos indicating that multiple donor gonadal germ cells formed functional oocytes in the two cohorts of surrogate host hens. A similar result was also observed for the two DDX4 ZW^-^ surrogate host females. We assayed the embryos for the presence of the iCaspase9 transgene carried by the female surrogate hosts. The 12 iCaspase9 females did not transmit the endogenous transgene suggesting that all embryos (634) were derived from donor gonadal germ cells. The ovaries from these hens showed a normal oocyte hierarchy (Supplementary Fig. 4A)

**Table 3.** Germline transmission from ♀ surrogate hosts injected with ♀ gonadal PGCs

| Host mating groups | No. of eggs laid per week * | Eggs set | Fertility<sup>§</sup> (% eggs set) | No. of RFP embryos (% fertile) | No. of GFP embryos (% fertile) | Transmission of surrogate iCaspase9 transgene (%) |
| --- | --- | --- | --- | --- | --- | --- |
| WT ♂ × 6 CRYO-2F♀ | 6.3 | 371 | 98% | 51 (14%) | 56 (36%) | 0 |
| WT ♂ × 6 CRYO-3F♀ | 5.0 | 340 | 80% | 66 (24%) | 150 (55%) | 0 |
| WT ♂ × 2 DDX4♀ | 6.4 | 148 | 93% | 28 (20%) | 49 (36%) | NA |

<sup>§</sup>Fertility was assessed between ED 4-7
\*Lay rate; eggs were counted over a 60 day period when hens were between 7-12 months of age and divided by the number of fertile hens present in pen. The maximum possible lay rate is 7.0 eggs per hen per week.

To demonstrate that we could produce pure offspring deriving from cryopreserved gonadal cells, we subsequently mated an individual iCaspase9 surrogate host male (CRYO1M_1179) to a cohort of iCaspase9 surrogate host females (CRYO2F) (Sire Dam Surrogate mating) (Table 4). The fertility of the laid eggs from the host hens ranged from 90-99% and the hatchability of the eggs was 91%. The presence of yellow (RFP^+^GFP^+^) offspring from this mating demonstrated that some chicks derived from cryopreserved donor male and donor female gonadal germ cells.

**Table 4.**
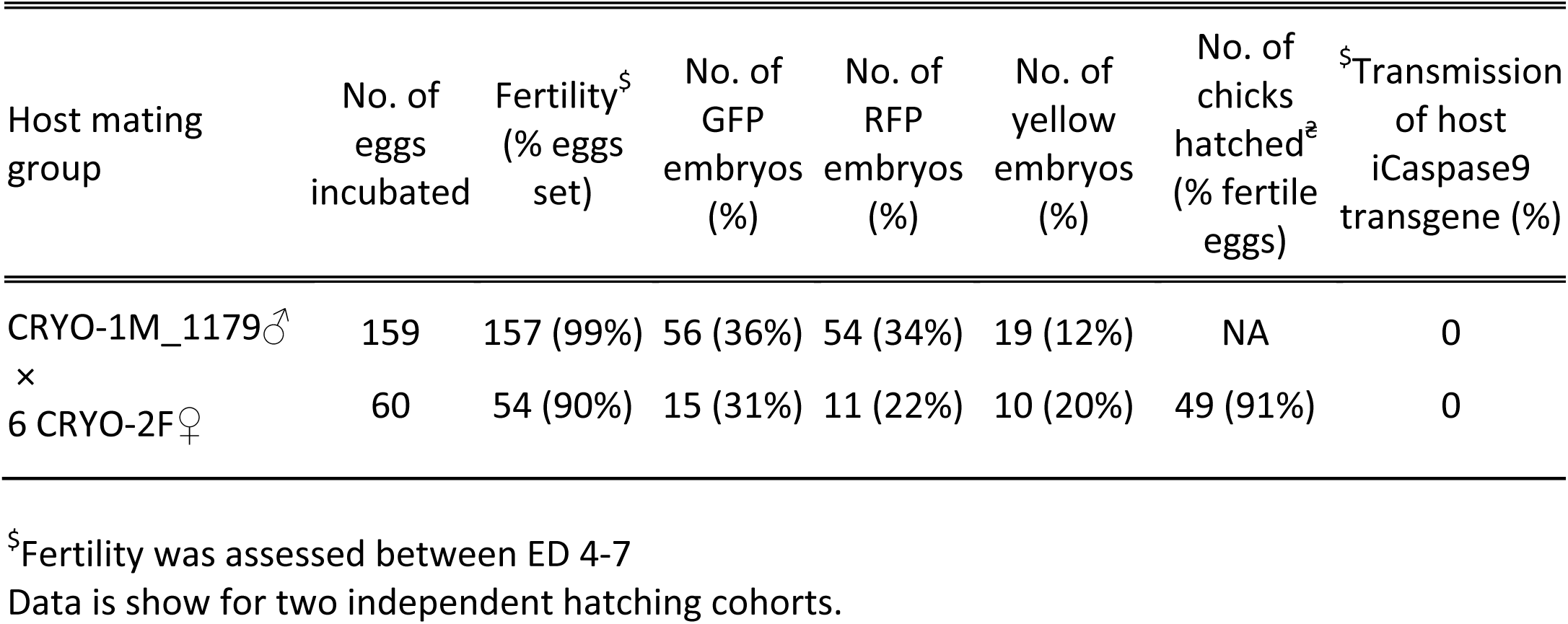
Fertility and hatching rate from ♂ surrogate hosts injected with ♂ gonadal cells mated to ♀ surrogate hosts injected with ♀ gonadal germ cells

### Donor germ cell transmission from opposite sex surrogate hosts

We have demonstrated previously that *in vitro* propagated chicken PGCs were not sex restricted for gamete formation and would produce functional gametes in opposite sex hosts, i.e., male PGCs in a female host differentiated into functional oocytes and female PGCs in a male surrogate host differentiated into functional spermatozoa (Ballantyne et al, in press). We now asked if gonadal donor cells carried by opposite sex iCaspase9 hosts produced functional gametes in the host testes or ovary. We observed that the iCaspase9 surrogate host hens injected with non-MACS-purified male gonadal germ cells (CRYO-1M) laid several (3) eggs. Examination of their ovaries after culling detected several mature yellow follicles in these hens (Supplementary Fig. 4B). We next asked if iCaspase9 host males carrying MACs-purified female germ cells generated functional spermatozoa. We naturally mated individual surrogate males carrying MACS-purified female gonadal germ cells (CRYO-3F) to wildtype females. We observed that fertility of the hens was over 95% and the occurrence of GFP^+^ embryos indicated that female gonadal germ cells were transmitted through the iCaspase9 host males (Table 5). Surprisingly, no RFP^+^ embryos were observed from this mating. Analysis of the testes of the iCapase9 host males revealed an unusual seminiferous tubule structure with a paucity of RFP^+^ cells on the abluminal surface and a reduction of differentiated spermatozoa in the luminal centre (Supplementary Fig. 3B and 5B). These results confirm that female gonadal germ cells can form functional spermatozoa in male hosts, achieve high levels of fertilisation (95-97%) and produce offspring.

**Table 5.**
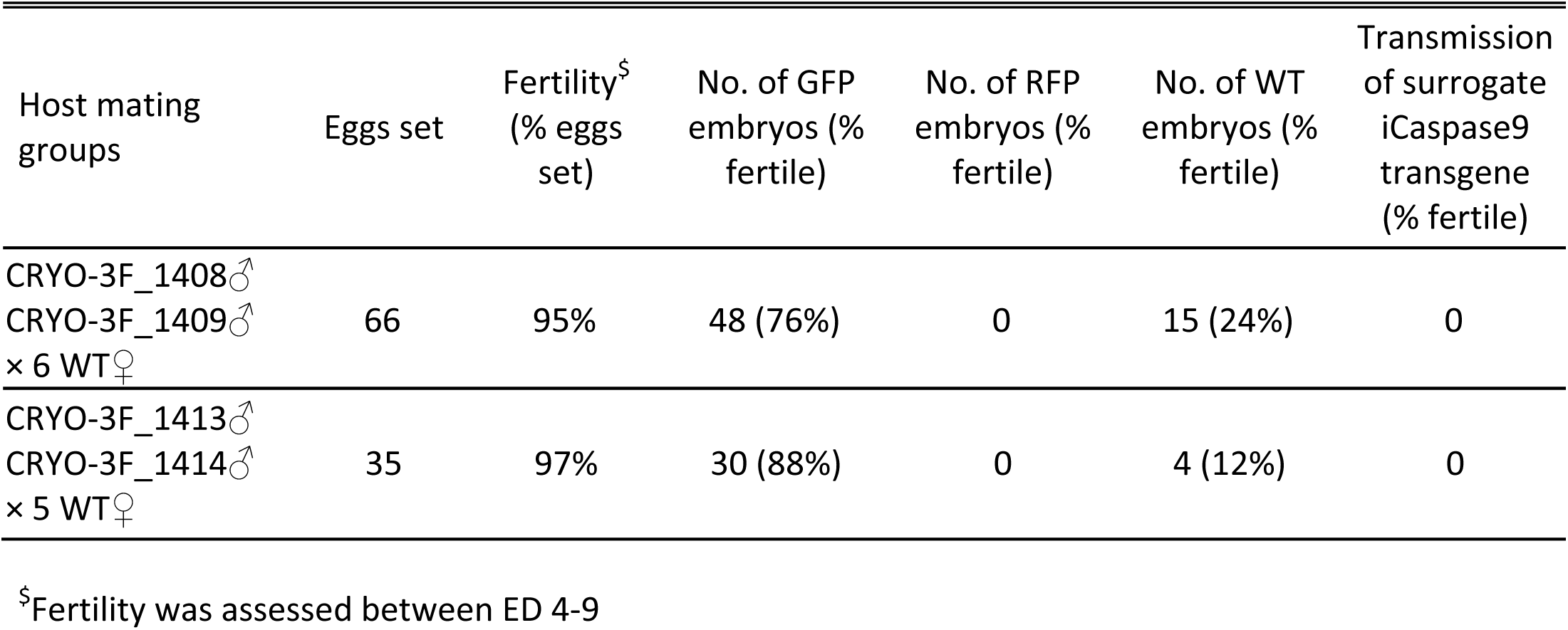
Germline transmission from ♂ surrogate hosts injected with ♀ gonadal germ cells

| Host mating groups | Eggs set | Fertility<sup>§</sup><br>(% eggs set) | No. of GFP embryos (% fertile) | No. of RFP embryos (% fertile) | No. of WT embryos (% fertile) | Transmission of surrogate iCaspase9 transgene (% fertile) |
| --- | --- | --- | --- | --- | --- | --- |
| CRYO-3F_1408♂<br>CRYO-3F_1409♂<br>× 6 WT♀ | 66 | 95% | 48 (76%) | 0 | 15 (24%) | 0 |
| CRYO-3F_1413♂<br>CRYO-3F_1414♂<br>× 5 WT♀ | 35 | 97% | 30 (88%) | 0 | 4 (12%) | 0 |

<sup>§</sup>Fertility was assessed between ED 4-9

### Transmission of multiple genotypes by surrogate hosts

The transmission of the RFP transgene was used as a proxy for measuring the transmission frequency of multiple genotypes through the surrogate hosts. The RFP^+^ donor gonads were from a heterozygote RFP^+^RFP^+^ male mated to heterozygote RFP^+^ females. Using this information, we calculated the expected number of RFP^+^ offspring if all donor genotypes were transmitted equally to the offspring (Table 6). For the first male surrogate host cohort, the donor gonadal material contained 10 GFP^+^: 2 RFP^+^ gonads. From this donor material we expected 11.1 % of the offspring to be RFP^+^ if each donor germ cell generated equal numbers of functional spermatozoa. For individual iCaspase9 males from one injection cohort (CRYO-1M), we observed 3%, 6%, and 28% of the embryos were RFP^+^; overall 14% of the embryos were RFP^+^. We then calculated if the transmission rates varied significantly from the expected values. RFP transmission of two of the three males differed significantly from the expected value, however the male cohort collectively did not vary significantly from the expected transmission rate. For the second cohort of males (CRYO-4M), the donor gonadal material contained 15 GFP^+^: 3 RFP^+^ gonads. From this donor material we again expected 11.1 % of the offspring to be RFP^+^. For individual iCaspase9 males from this cohort we observed that 5%, 5%, 12%, and 21% of the embryos were RFP^+^ and overall, 11% of the embryos were RFP^+^. Three of the four males diverged significantly from the expected transmission rate but again the male cohort considered collectively did not vary significantly from the expected transmission rate. These data indicate that individual males transmitted donor germ cell genotypes at varying frequencies. However, combined data from multiple host iCaspase9 males from a single injection cohort suggested that all donor genotypes were transmitted proportionally to the offspring.

**Table 6.** RFP transmission rates identify multiple transmission events

| Bird ID | total eggs set | Fertile eggs<sup>§</sup> (%) | No. of GFP (%) | No. of RFP (%) | No. of WT (%) | No. of iCaspase9 (%) | No. of gonad pairs | expected RFP (%) | p value |
| --- | --- | --- | --- | --- | --- | --- | --- | --- | --- |
| CRYO-1M: wb1181 | 147 | 141<br>96% | 101<br>72% | 4<br>3% | 36<br>26% | 0 | 10 GFP<br>+2 RFP | 11.1% | <0.001 |
| CRYO-1M: wb1182 | 148 | 147<br>99% | 89<br>61% | 9<br>6% | 48<br>33% | 9<br>6% | 10 GFP<br>+2 RFP | 11.1% | 0.05 |
| CRYO-1M: wb1179 | 187 | 184<br>98% | 55<br>30% | 51<br>28% | 76<br>41% | 0 | 10 GFP<br>+2 RFP | 11.1% | <0.001 |
| total | 482 | 472<br>98% | 245<br>52% | 64<br>14% | 160<br>34% | 9<br>2% | 10 GFP<br>+2 RFP | 11.1% | 0.106 |
| CRYO-2F | 371 | 362<br>98% | 137<br>38% | 51<br>14% | 174<br>48% | 0 | 10 GFP<br>+4 RFP | 19.0% | 0.016 |
| CRYO-3F | 340 | 272<br>80% | 150<br>55% | 66<br>24% | 55<br>20% | 0 | 17 GFP<br>+6 RFP | 17.4% | 0.005 |
| CRYO-4M: wb1420 | 89 | 81<br>91% | 64<br>79% | 4<br>5% | 13<br>16% | 0 | 15 GFP<br>+3 RFP | 11.1% | 0.078 |
| CRYO-4M: wb1424 | 60 | 56<br>93% | 41<br>73% | 3<br>5% | 11<br>20% | 0 | 15 GFP<br>+3 RFP | 11.1% | 0.16 |
| CRYO-4M: wb1427 | 74 | 63<br>85% | 29<br>46% | 13<br>21% | 21<br>33% | 0 | 15 GFP<br>+3 RFP | 11.1% | 0.043 |
| CRYO-4M: wb1430 | 143 | 118<br>83% | 80<br>68% | 14<br>12% | 24<br>20% | 0 | 15 GFP<br>+3 RFP | 11.1% | 0.884 |
| total | 366 | 318<br>87% | 214<br>67% | 34<br>11% | 69<br>22% | 0 | 15 GFP<br>+3 RFP | 11.1% | 0.859 |
| CRYO-5F | 148 | 138<br>93% | 49<br>36% | 28<br>20% | 58<br>42% | 0 | 15 GFP<br>+3 RFP | 11.1% | 0.003 |

<sup>§</sup>Fertility was measured for between ED 4-6.
A statistical analysis was performed to determine if the no. of observed RFP<sup>+</sup> embryos differed significantly from the no. of expected RFP<sup>+</sup> embryos. A p value of <0.05 was designated as the value at which the observed and expected numbers differed significantly.

For the first iCaspase9 female host cohort, which were housed collectively, the donor gonadal material contained 10 GFP^+^: 4 RFP^+^ gonads. From this donor material we expected that 19.0% of the offspring would be RFP^+^ if each donor germ cell generated equal numbers of functional ova. We observed that the overall transmission rate for the cohort was 14% which did not deviate significantly from the expected transmission rate (Table 6). For the second iCaspase9 cohort of females, the donor gonadal material was mixed 17 GFP^+^: 6 RFP^+^ gonads. From this donor material we expected that 17.4% of the offspring would be RFP^+^ if each donor germ cell generated equal numbers of functional ova. We observed that the overall transmission rate was 24% which did not deviate significantly from the expected transmission rate.

For the DDX4 surrogate host females, we expected that 11.1% of the offspring would be RFP^+^. We observed that 20% of the offspring were RFP^+^. This value differed significantly from the expected transmission rate. These results show that the iCaspase9 females transmitted multiple genotypes at expected rates whereas the small cohort of DDX4 females (2) did not. Overall, these data suggest that donor gonadal germ cells deriving from different genotypes will transmit with equal efficacy through same sex male and female iCapsase9 surrogate hosts.

## Discussion

Here we demonstrate a method to simply and efficiently cryopreserve reproductive embryonic gonads from the chicken. Multiple samples can be quickly dissected, visually sexed, pooled, and cryopreserved to provide a frozen genetic resource for chicken breeds. This cryopreservation method will allow biobanking of poultry breeds to be carried out in localities lacking extensive infrastructure and equipment. Re-establishing poultry breeds from frozen material remains technically demanding. Breed regeneration, however, can be separated spatially and temporally from the storage facilities, i.e. the biobank. We chose to purify female germ cells using MACS, in place of florescent activated cell sorter (FACS), as MACS will be more applicable in LMICs. It remains to be tested if MACS enrichment is necessary to achieve germ line transmission from female gonadal cells. We expect that MACS enrichment of male gonadal cells would achieve functional gametogenesis in female iCapase9 hosts as we observed that several eggs were laid when using male dissociated gonadal cells. Germline transmission from gonadal cells in opposite sex hosts may be useful for increasing overall genetic diversity of the regenerated flock as long as inbreeding is avoided. This technology can also be applied immediately at chicken research facilities in most high income countries (HICs). Our GFP and RFP chicken lines carrying a used in this demonstration contain a single transgene insert (McGrew, Sherman et al. 2008, Ho, Freem et al. 2019). Robust numbers of offspring were generated from frozen gonadal material for both sexes. To cryopreserving transgenic reporter lines of chicken, it would be simpler and more efficient to store multiple vials of male gonads for future injections.

Local indigenous breeds commonly consist of small populations with low egg production (Dessie and Ogle 2001, Melesse 2014). Here, we used 12-23 donor embryos per injection experiment, but 6-7 donor embryo gonads should be sufficient to generate an injection cohort of surrogate hosts (Table 1). This number of eggs could be obtained from small flocks of indigenous chicken. Further research is needed to determine if donor germ cell genotypes are transmitted equally to the offspring or if some genotypes are underrepresented or overrepresented in the offspring. Our data show that mixing GFP: RFP donor gonadal cells at a 5:1 ratio maintained this ratio, on average, in the offspring of the iCapase9 host birds. In our experiments, however, the genetics of the donor gonadal material (commercial brown layer) matched the genetics of the iCaspase9 and *DDX4* surrogate host birds (commercial brown layer). It remains to be shown if donor gonadal material from rare breeds and indigenous chicken ecotypes-many with poor laying and smaller populations-will transmit donor genotypes equally in the iCaspase9 brown layer host. This question can be addressed by genotyping both the donor material and the surrogate host offspring to measure the transmission frequencies of multiple genotypes. In animal species, it is hypothesised that sperm competition occurs when multiple males mate individual females (Amann, Saacke et al. 2018, Birkhead and Montgomerie 2020). In this case, we may observe that donor germ cells of different genotypes do not compete equally during gametogenesis in an individual host cockerel.

Chicken flocks are highly susceptible to loss of fitness from inbreeding which leads to poor gamete quality resulting in low fertilizing potential, reduced egg laying and diminished hatchability. Regenerated chicken flocks must therefore consist of numerous genotypes in order to avoid genetic bottlenecks and the accompanying reductions in reproductive fitness of the flock. FAO advises that to maintain rare and local livestock breed populations that suffer from minimal inbreeding (no greater than 1%), a population of 13 unrelated males bred to 13 unrelated females (from independent family groups) would be needed to revive a population (FAO 1998). Here, in our exemplar, we mated single surrogate host males carrying 12 donor genotypes to a cohort of surrogate host females carrying 14 donor genotypes. This number of genotypes would theoretically regenerate a genetically diverse population with an inbreeding coefficient less than 1% and meet FAO guidelines.

The methodology presented here relies on sterile surrogate hosts for efficient breed regeneration. We generated greater numbers of donor/host sex-matched chimeras using the iCapsase9 transgenic chicken host than the *DDX4* knockout female host. The *DDX4* female host, however, cannot transmit the endogenous transgene to its offspring (Taylor, Carlson et al. 2017, Woodcock, Gheyas et al. 2019). In contrast, in the iCaspase9 host, 1/7 males and 0/12 female hosts generated transgenic offspring. Genetic modifications of the iCapsase9 transgene should improve the induced sterility by B/B compound of surrogate host embryos. The use of a genetically modified or genome edited surrogate host chicken for biobanking platforms will require that new livestock regulations are adopted in the countries implementing this technology. In the future, we envision that improved non-GM sterility protocols could replace the current surrogate hosts and eliminate the need for the development of new regulations (Macdonald, Glover et al. 2010, Nakamura, Usui et al. 2010, Nakamura, Usui et al. 2012). Chemical and physical sterility treatments, however, have a high impact on the health of the host bird. In contrast, genome editing loss of function mutations and iCapase9 induced apoptosis has minimal to no welfare impact on the surrogate host birds. As the use of GM sterile surrogate hosts is more sustainable and scalable and supports the principles of the 3Rs, it might be the ideal approach provided that it could be ensured that no endogenous germ cell transmission occurred. Nevertheless, as future research improves surrogate host technology, the cryopreserved chicken gonadal tissues will remain a functional resource to secure biodiversity and sustainability for future poultry farming.

## Materials and Methods

### Chicken breeds and embryos

Fertile eggs from TdTomato transgenic chickens (ubiquitous expression of TdTomato (RFP^+^)(Ho, Freem et al. 2019)), GFP^+^ transgenic chickens (ubiquitous expression of GFP (McGrew, Sherman et al. 2008)) and Hy-line Brown layer were obtained from NARF at the Roslin Institute (https://www.ed.ac.uk/roslin/national-avian-research-facility). The breeding of TdTomato transgenic chickens were a heterozygous x heterozygous cross, only the tissues from RFP embryos were dissected for cryopreservation. The breeding of GFP transgenic chicken included both heterozygous x heterozygous crosses and homozygous x wild type crosses. The Caspase9 line of chickens were generated using a Hy-line Brown layer PGCs. Heterozygous and homozygous cockerels carrying the iCaspase9 transgene were crossed to Hy-line hens to produce fertile eggs for injection and hatching. All three lines were maintained on a Hy-line Brown background. The fertile eggs were incubated at 37.8°C under humid conditions with rocking. Embryonic development was staged according to the morphological criteria of Hamburger and Hamilton. Stage 35 (day 9) was principally decided by eye morphology. All animal management, maintenance and embryo manipulations were carried out under UK Home Office license and regulations. Experimental protocols and studies were approved by the Roslin Institute Animal Welfare and Ethical Review Board Committee.

### Isolation and cryopreservation of gonads

Embryos between ED 9-12 days of incubation were used for the isolation of gonadal tissues. In a clean laminar flow hood, after wiping surface of eggs with 70% ethanol the blunt end of the egg shell was cracked open using forceps and the shell membrane removed to visualize the embryo. The embryo was isolated and placed in a 100mm petri dish, effectively culling the embryo by severing the neck to decapitate the embryo using forceps or scissors. Under a stereo microscope, the embryo body was positioned by placing the ventral surface (belly) upwards, the embryo was opened and the visceral organs carefully removed to expose the gonads and the attached mesonephroi. The embryo was visually sexed by gonadal morphology. Males have two elliptical, ‘sausage-shaped’ gonads of approximately equal sizes. Females have a much larger left gonad that is flattened, a ‘pancake-shape’. Both gonads were gently dissected off the mesonephroi using a 23G (1 ¼’’ in length) hypodermic needle. The dissected gonads were picked up using the needle tip and transferred into a drop of DMEM on the peripheral area of the dissection petri dish, to wash off blood cells. The gonad tissues, 5 pairs of GFP and 1 pair of RFP for each sex, were then transferred into a 1.5 ml eppendorf tube (screw top) containing 500 μl cold DMEM (separate tubes for each sex) and placed on ice.

To cryopreserve the material, the gonads were moved to the tube bottom by a quick 4 second spin of the eppendorf tubes in a benchtop centrifuge. The DMEM medium on top was gently removed. 100 μl Stem-Cellbanker were added to the tubes for a first medium exchange. After another quick spin to re-collect the gonads at tube bottom, supernatant on top was gently removed and 200 μl Stem-Cellbanker was added to the tissues. After 15 min equilibration of gonadal tissues in Stem-Cellbanker on ice, the tubes were placed into a Mr. Frosty™ Freezing Container and placed in a -80°C freezer overnight, and then transferred into a -150°C freezer or liquid nitrogen for long-term storage.

### Single gonadal cell preparation and magnetic-activated cell sorting (MACS) using SSEA-1 antibody

The gonadal tissues were retrieved from the -150°C freezer and thawed at 37°C for 30 sec in a heating block. In a bio-safety hood, the Stem-Cellbanker was gently removed from tubes and 500 μl DMEM was added slowly drop wise to wash and equilibrate the tissues. The tissues were re-collected at the tube bottom by a quick spin. After removing the wash solution, 200 μl dispase/collagenase solution (5 mg/ml in PBS) was added to the tissues. The tubes were incubated at 37°C for 10 min to dissociate the gonads, shaking the tubes to resuspend the tissues three times during the incubation period. Single cells were released by triturating the tissues up and down using a P200 pipette until tissue clumps disappeared. The cell suspension was filtered through a 40 μm pore cell strainer, followed by washing the cell strainer with MACS sorting buffer (0.5% BSA, 2 mM EDTA in PBS) three times, 1ml each time. The filtered cell suspension was aliquoted into 1.5 ml eppendorf tubes and centrifuged at 2000 rpm for 4 min to pellet the cells. The cells were washed twice with 500 μl DMEM then the cell number was determined using a haemocytometer. The final cell density was adjusted by adding additional DMEM. For male cells, cell density was adjusted to 50,000 – 60,000 cells/ μl for subsequent injection of surrogate hosts.

The dissociated female gonadal cells were enriched by MACS using an SSEA-1 antibody before injection. The female cells were resuspended in MACS buffer to a cell density of 2 × 10^7^ cells/ml (∼200 ul). SSEA1 antibody was added to the cell suspension at 1.5 μg antibody per 10^7^ cells. The solution was incubated on a roller for 20 min at 4°C. The solution was centrifuged at 2000 rpm for 4 min to pellet the cells. The cells were washed twice with 200 μl MACS buffer, and the cells were resuspended in appropriate volume of MACS buffer to reach a cell density of 1 × 10^8^ cells/ml (∼40 μl). Anti-mouse IgM-conjugated MACS beads (∼10 μl) were added (2.0 μl of beads per 10^7^ cells). The cell solution was incubated on a roller for 20 min at 4°C. MACS buffer was added to a final volume of 1 ml, and centrifuged at 2000 rpm for 4 min to separate the conjugated cells from the excess microbeads. The cell pellet was resuspended in 1 ml of MACS buffer and loaded into a LS column mounted onto a magnetic station. The column was washed with 3 ml MACS buffer twice, the column was taken off from the magnetic station and the bound cells were eluted with 3 ml MACS buffer into a 15 ml falcon tube. The cells were pelleted by centrifuge at 2000 rpm for 4 min. After washing the cells twice with 200 μl DMEM, the cells were resuspended in DMEM at a cell density of 1,000 cells/ul and injected into host embryos.

### Flowcytometric analysis

The single gonadal cells were immunostained by SSEA-1 antibody (1:500 dilution) for 20 min on ice, followed by anti-mouse IgM conjugated by AF654 (1:5000 dilution) for 15 min on ice. The resulted staining was analysed by B&D Fortessa and Flowjo v10 software.

### Colonisation experiments

Female gonads from ED 8 to 11 GFP^+^ and RFP^+^ transgenic embryos were pooled and frozen in Stem-Cellbanker at -150°C. The single cells were thawed and dissociated as above using dispase/collagenase enzyme and 15,000 cells, except of 10,000 cells/embryos for E9 donor cells, were injected into non-fluorescent host embryos at ED 2.5 (stage 16 HH). The migration and colonization of donor cells in surrogate host were observed at ED 8 or 9. The pools for donor female gonadal tissues were 6 GFP^+^: 1 RFP^+^ for ED 8, 6 GFP^+^: 2 RFP^+^ for ED 9, 7 GFP^+^ for ED 10 and 3 GFP^+^: 1 RFP^+^ male for ED 11.

For hatching surrogate hosts, ED 2.5 iCaspase9 host embryos were injected directly with male gonadal donor cells mixed with B/B compound. Female donor gonadal cells were MAC-sorted before mixing with B/B compound and injected. A 1.0 μl cell solution containing 0.5 mM f.c. AP20187 (B/B) compound (Takara Bioscience) was injected into the dorsal aorta through a small window made in the ventral egg. 50 μl of Penicillin/Streptomycin (P/S) solution (containing 15 uM f.c. B/B compound) was pipetted on top of the embryo before sealing. The window was sealed with a 0.5 cm square of Leukosilk (BSN Medical) and the injected eggs were incubated until hatching. Hatchlings were raised until sexual maturity when natural matings in floor pens were set up.

### Statistics

The expected proportion of RFP chick in offspring could be predicted by RFP allele frequency in a pool of cryopreserved gonadal tissues, using formula ((4/3) x number of RFP gonad pairs) /(2 x total number of gonad pairs in a pool). The RFP germline transmission frequency was statistically analysed by comparison of the expected and observed RFP percentages by using the exact one proportion function in basic statistics of Minitab. Other statistical analyses were calculated using a two-tail student *t* test. The error bars in all figures are S.E.M.

## Acknowledgements

We thank Norman Russell for the chicken photographs and Helen Brown for statistical advice. We thank Phil Purdy for commenting on the manuscript. We thank the staff at the National Avian Research Facility (NARF) for care and maintenance of the birds used in this study. This work was supported by the Institute Strategic Grant Funding from the BBSRC (BB/P0.13732/1 and BB/P013759/1) and the NC3Rs (C/V001124/1). Primary funding of this research was from the Bill & Melinda Gates Foundation and with UK aid from the UK Foreign, Commonwealth and Development Office (Grant Agreement OPP1127286) under the auspices of the Centre for Tropical Livestock Genetics and Health (CTLGH), established jointly by the University of Edinburgh, SRUC (Scotland’s Rural College), and the International Livestock Research Institute. The findings and conclusions contained within are those of the authors and do not necessarily reflect positions or policies of the Bill & Melinda Gates Foundation nor the UK Government.

## Supplementary Figures

**Supplementary Fig. 1.**
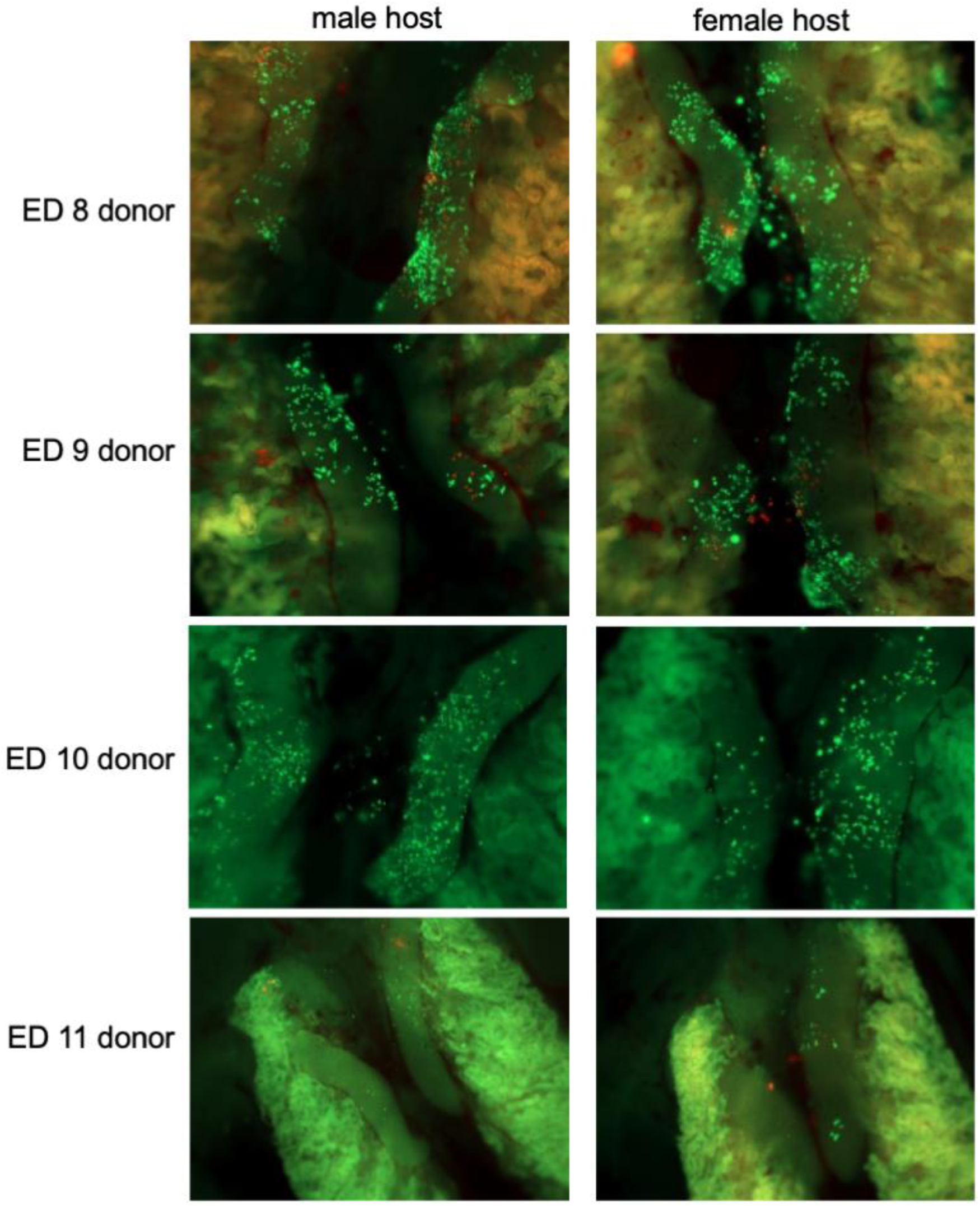
Recolonization of surrogate host embryos by frozen gonadal germ cells Gonadal cell suspensions from cryopreserved ED 8 to 11 GFP^+^ and RFP^+^ female embryos (5 GFP^+^:1 RFP^+^) were injected into wildtype ED 2.5 (stage 16 HH) host embryos. Host embryos were examined at ED 8-9 of incubation for gonadal colonisation. 15,000 cells were injected per embryo (10,000 cells/embryos for ED 9 donor cells). The representative images are from at least 3 independent injection experiments.

**Supplementary Fig. 2.**
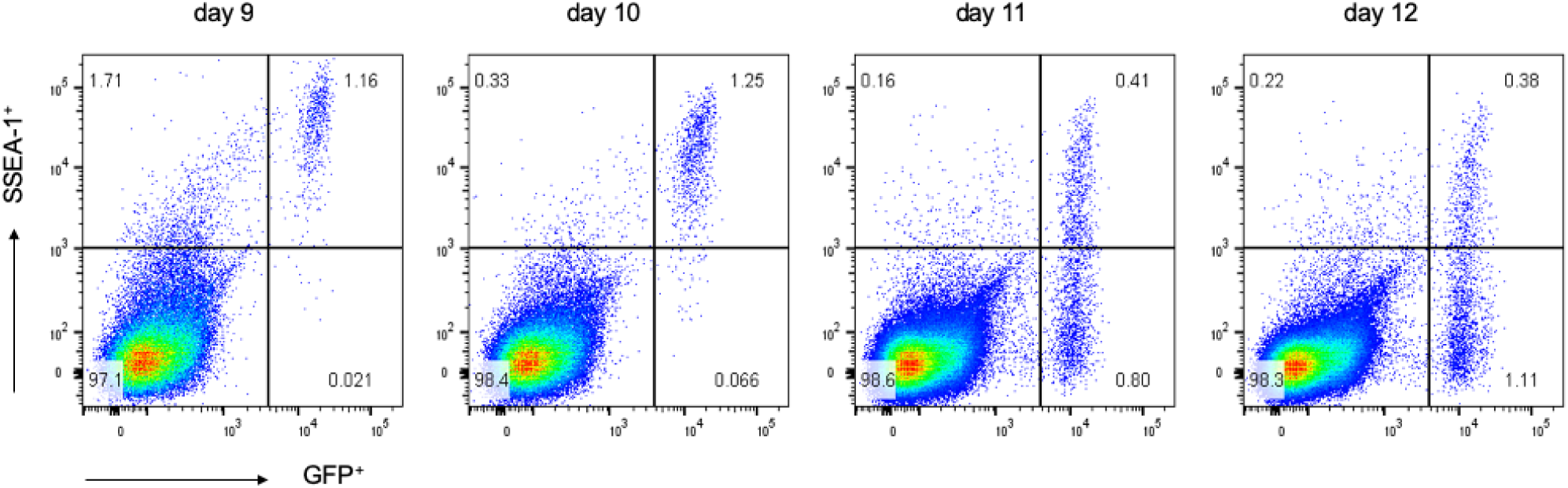
SSEA-1 expression of post-migratory gonadal germ cells iCaspase9 female embryos were used to identify putative germ cells using GFP expression. Gonadal cells were dissociated and examined by flow cytometry for GFP and SSEA-1 expression. SSEA-1 co-expression by GFP^+^ cells: 98.2% at day 9, 94.7 at day 10, 51.3% at day 11, and 34.2% at day 12. n=4-8 embryos independently assayed for each time point.

**Supplementary Fig. 3.**
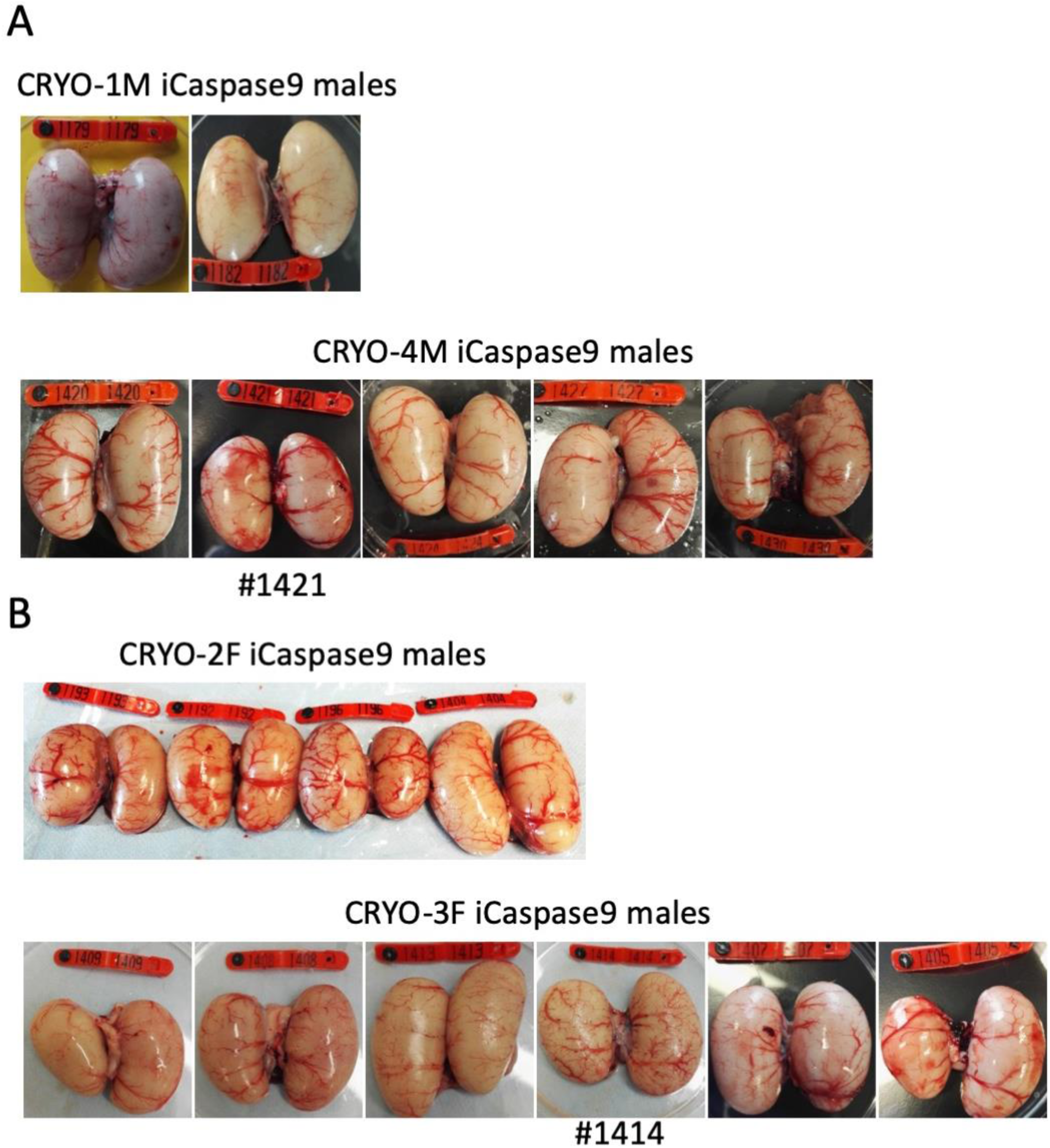
Adult gonads of iCaspase9 surrogate host cockerels A. Male surrogate hosts injected with male gonadal germ cells examined at >25 weeks of age. B. Male surrogate hosts injected with female gonadal germ cells examined at >25 weeks of age. Red wing tags = 5 cm.

**Supplementary Fig. 4.**
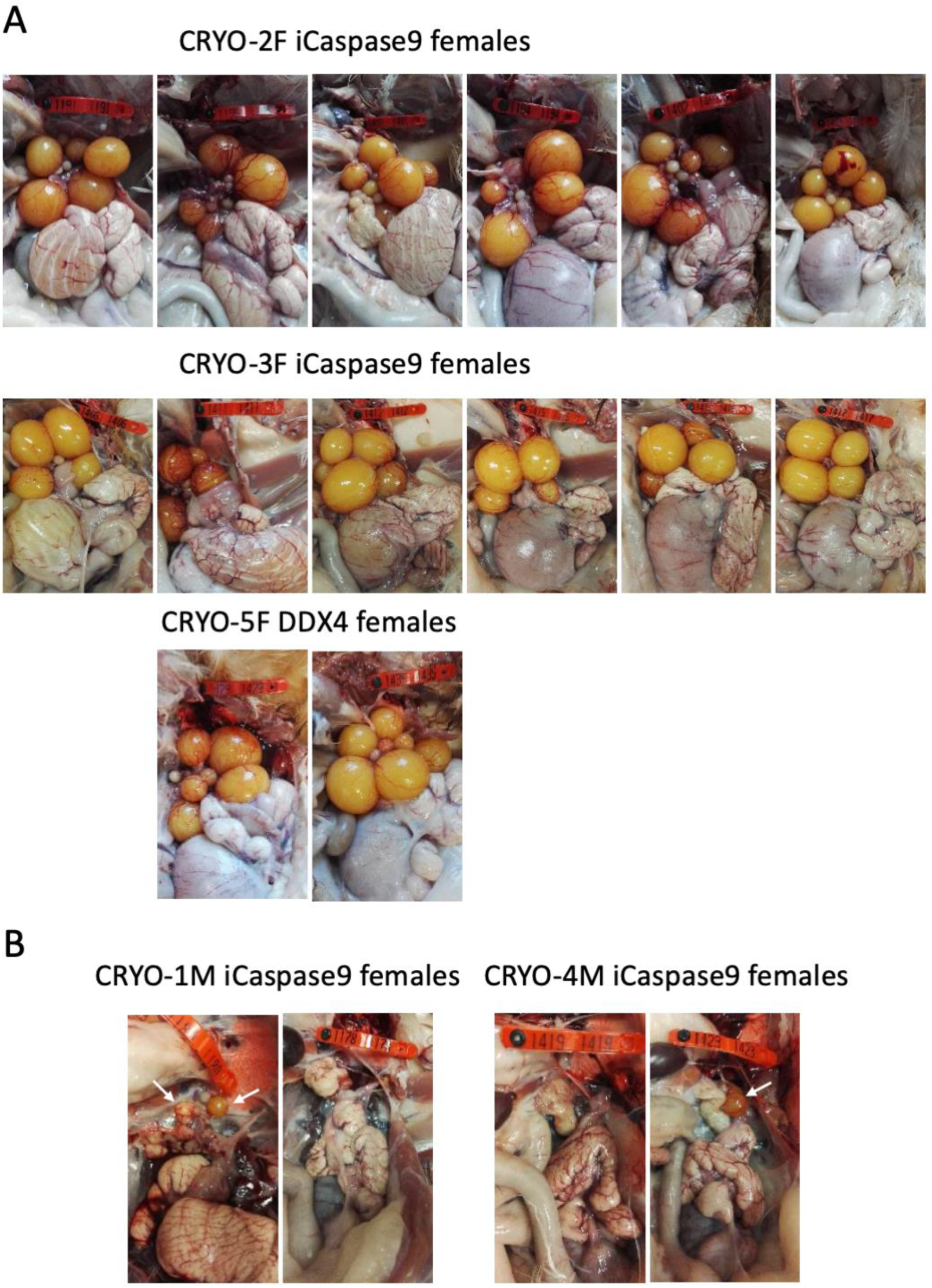
Adult gonads of iCaspase9 surrogate host hens A. Female surrogate hosts injected with female gonadal germ c3ells examined at >25 weeks of age. B. Female surrogate hosts injected with male gonadal germ cells examined at >25 weeks of age. White arrows, yellow follicles. Red wing tags = 5 cm.

**Supplementary Fig. 5.**
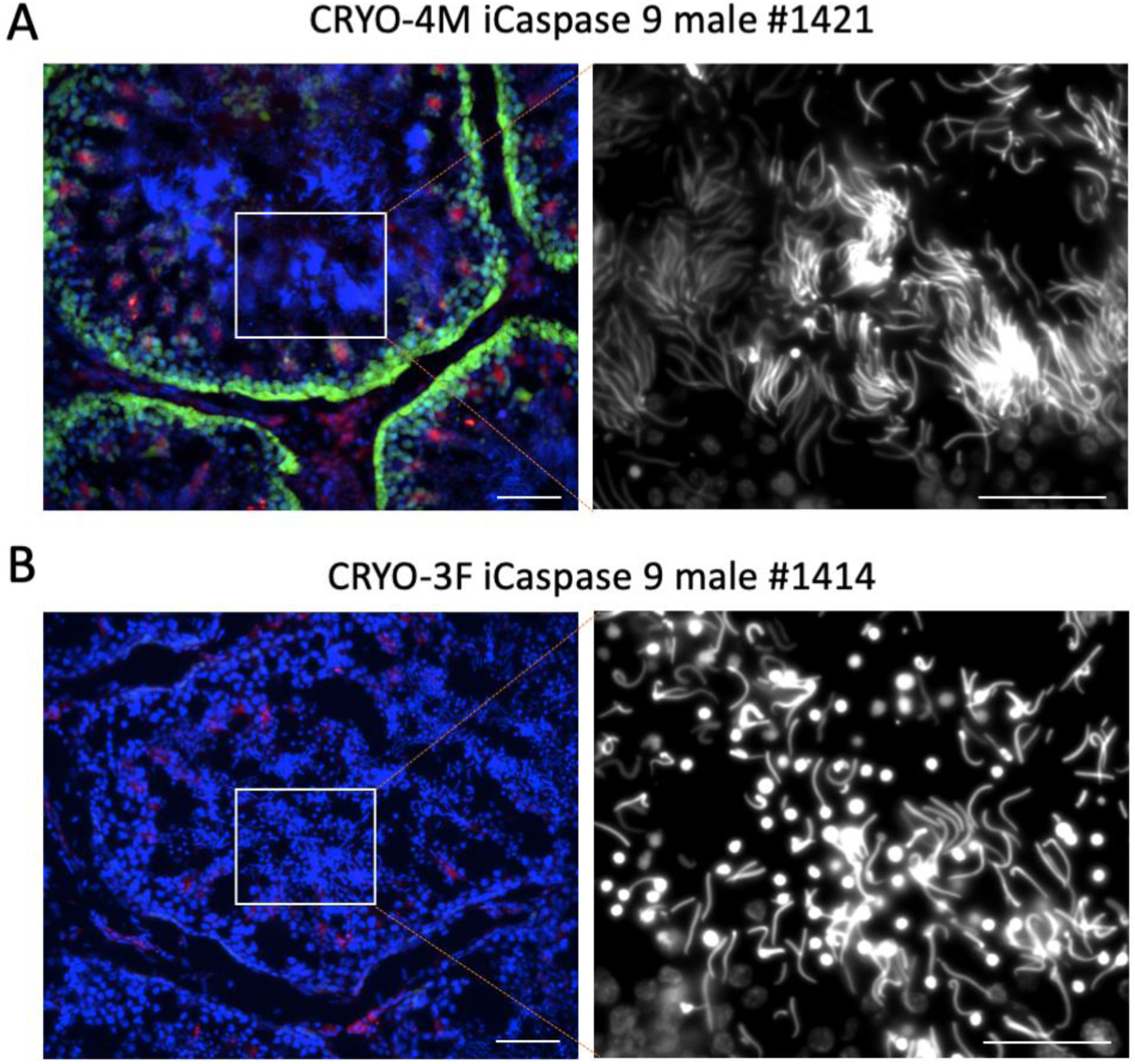
Cryosections of adult testes of iCaspase9 surrogate host cockerels A. A male surrogate host injected with male gonadal germ cells examined at >25 weeks of age. Blue, Hoechst stain; right panel, Hoechst stain shown in white. *, lumen of tubule B. Male surrogate hosts injected with female gonadal germ cells examined at >25 weeks of age. Blue, Hoechst stain; right panel, Hoechst stain shown in white. Scale bars = 50 um.

## References

Alders, R. G. and R. A. E. Pym (2009). “Village poultry: still important to millions, eight thousand years after domestication.” Worlds Poultry Science Journal 65(2): 181–190.

Amann, R. P., R. G. Saacke, G. F. Barbato and D. Waberski (2018). “Measuring Male-to-Male Differences in Fertility or Effects of Semen Treatments.” Annual Review of Animal Biosciences 6(1): 255–286.

Ballantyne, M., M. Woodcock, D. Doddamani, T. Hu, L. Taylor, R. J. Hawken and M. J. McGrew (2021). “Direct allele introgression into pure chicken breeds using Sire Dam Surrogate (SDS) mating.” Nat Commun 12(1): 659.

Birkhead, T. R. and R. Montgomerie (2020). “Three decades of sperm competition in birds.” Philos Trans R Soc Lond B Biol Sci 375(1813): 20200208.

Blesbois, E., I. Grasseau, F. Seigneurin, S. Mignon-Grasteau, M. Saint Jalme and M. M. Mialon-Richard (2008). “Predictors of success of semen cryopreservation in chickens.” Theriogenology 69(2): 252–261.

DAD-IS (2021). FAO.

Davey, M. G., A. Balic, J. Rainger, H. M. Sang and M. J. McGrew (2018). “Illuminating the chicken model through genetic modification.” Int J Dev Biol 62(1-2-3): 257–264.

Dessie, T. and B. Ogle (2001). “Village Poultry Production Systems in the Central Highlands of Ethiopia.” Tropical Animal Health and Production 33(6): 521–537.

FAO (1998). Secondary guidelines: Management of small populations at risk. FAO: 1–210.

Fulton, J. E. and M. E. Delany (2003). “Genetics. Poultry genetic resources--operation rescue needed.” Science 300(5626): 1667–1668.

Ho, W. K. W., L. Freem, D. Zhao, K. J. Painter, T. E. Woolley, E. A. Gaffney, M. J. McGrew, A. Tzika, M. C. Milinkovitch, P. Schneider, A. Drusko, F. Matthaus, J. D. Glover, K. L. Wells, J. A. Johansson, M. G. Davey, H. M. Sang, M. Clinton and D. J. Headon (2019). “Feather arrays are patterned by interacting signalling and cell density waves.” PLoS Biol 17(2): e3000132.

Hughes, G. C. (1963). “The Population of Germ Cells in the Developing Female Chick.” J Embryol Exp Morphol 11: 513–536.

Karagenc, L., Y. Cinnamon, M. Ginsburg and J. N. Petitte (1996). “Origin of primordial germ cells in the prestreak chick embryo.” Dev Genet 19(4): 290–301.

Kim, J. N., M. A. Kim, T. S. Park, D. K. Kim, H. J. Park, T. Ono, J. M. Lim and J. Y. Han (2004). “Enriched gonadal migration of donor-derived gonadal primordial germ cells by immunomagnetic cell sorting in birds.” Mol Reprod Dev 68(1): 81–87.

Macdonald, J., J. D. Glover, L. Taylor, H. M. Sang and M. J. McGrew (2010). “Characterisation and Germline Transmission of Cultured Avian Primordial Germ Cells.” Plos One 5(11).

Matsuzaki, M., N. Hirohashi, S. Mizushima and T. Sasanami (2021). “Effect of sperm surface oligosaccharides in sperm passage into sperm storage tubules in Japanese quail (Coturnix japonica).” Anim Reprod Sci 227: 106731.

McGrew, M. J., A. Sherman, S. G. Lillico, F. M. Ellard, P. A. Radcliffe, H. J. Gilhooley, K. A. Mitrophanous, N. Cambray, V. Wilson and H. Sang (2008). “Localised axial progenitor cell populations in the avian tail bud are not committed to a posterior Hox identity.” Development 135(13): 2289–2299.

Melesse, A. (2014). “Significance of scavenging chicken production in the rural community of Africa for enhanced food security.” Worlds Poultry Science Journal 70(3): 593–606.

Mozdziak, P. E., J. Angerman-Stewart, B. Rushton, S. L. Pardue and J. N. Petitte (2005). “Isolation of chicken primordial germ cells using fluorescence-activated cell sorting.” Poult Sci 84(4): 594–600.

Mozdziak, P. E., R. Wysocki, J. Angerman-Stewart, S. L. Pardue and J. N. Petitte (2006). “Production of chick germline chimeras from fluorescence-activated cell-sorted gonocytes.” Poult Sci 85(10): 1764–1768.

Naito, M., T. Minematsu, T. Harumi and T. Kuwana (2007). “Testicular and ovarian gonocytes from 20-day incubated chicken embryos contribute to germline lineage after transfer into bloodstream of recipient embryos.” Reproduction 134(4): 577–584.

Nakamura, Y., F. Usui, D. Miyahara, T. Mori, T. Ono, H. Kagami, K. Takeda, K. Nirasawa and T. Tagami (2012). “X-irradiation removes endogenous primordial germ cells (PGCs) and increases germline transmission of donor PGCs in chimeric chickens.” J Reprod Dev 58(4): 432–437.

Nakamura, Y., F. Usui, T. Ono, K. Takeda, K. Nirasawa, H. Kagami and T. Tagami (2010). “Germline replacement by transfer of primordial germ cells into partially sterilized embryos in the chicken.” Biol Reprod 83(1): 130–137.

Nakamura, Y., Y. Yamamoto, F. Usui, Y. Atsumi, Y. Ito, T. Ono, K. Takeda, K. Nirasawa, H. Kagami and T. Tagami (2008). “Increased proportion of donor primordial germ cells in chimeric gonads by sterilisation of recipient embryos using busulfan sustained-release emulsion in chickens.” Reprod Fertil Dev 20(8): 900–907.

Ritchie, H. and M. Roser. (2021). Retrieved March 19, 2021, from https://ourworldindata.org/meat-production#number-of-animals-slaughtered.

Tajima, A., M. Naito, Y. Yasuda and T. Kuwana (1998). “Production of germ-line chimeras by transfer of cryopreserved gonadal primordial germ cells (gPGCs) in chicken.” Journal of Experimental Zoology 280(3): 265–267.

Taylor, L., D. F. Carlson, S. Nandi, A. Sherman, S. C. Fahrenkrug and M. J. McGrew (2017). “Efficient TALEN-mediated gene targeting of chicken primordial germ cells.” Development 144(5): 928–934.

Thelie, A., A. Bailliard, F. Seigneurin, T. Zerjal, M. Tixier-Boichard and E. Blesbois (2019). “Chicken semen cryopreservation and use for the restoration of rare genetic resources.” Poult Sci 98(1): 447–455.

Urven, L. E., C. A. Erickson, U. K. Abbott and J. R. McCarrey (1988). “Analysis of germ line development in the chick embryo using an anti-mouse EC cell antibody.” Development 103(2): 299–304.

van de Lavoir, M. C., C. Mather-Love, P. Leighton, J. H. Diamond, B. S. Heyer, R. Roberts, L. Zhu, P. Winters-Digiacinto, A. Kerchner, T. Gessaro, S. Swanberg, M. E. Delany and R. J. Etches (2006). “High-grade transgenic somatic chimeras from chicken embryonic stem cells.” Mech Dev 123(1): 31–41.

Whyte, J., J. D. Glover, M. Woodcock, J. Brzeszczynska, L. Taylor, A. Sherman, P. Kaiser and M. J. McGrew (2015). “FGF, Insulin, and SMAD Signaling Cooperate for Avian Primordial Germ Cell Self-Renewal.” Stem cell Reports 5(6): 1171–1182.

Whyte J. B. E., Mcgrew M.J. (2015). Increased Sustainability in Poultry Production: New Tools and Resources for Genetic Management. CABI Publishing, CABI Publishing.

Woelders, H., J. Windig and S. Hiemstra (2012). “How Developments in Cryobiology, Reproductive Technologies and Conservation Genomics Could Shape Gene Banking Strategies for (Farm) Animals.” Reproduction in Domestic Animals 47(s4): 264-273.

Woodcock, M. E., A. A. Gheyas, A. S. Mason, S. Nandi, L. Taylor, A. Sherman, J. Smith, D. W. Burt, R. Hawken and M. J. McGrew (2019). “Reviving rare chicken breeds using genetically engineered sterility in surrogate host birds.” Proc Natl Acad Sci U S A 116(42): 20930–20937.

Yang, S. Y., H. J. Lee, H. C. Lee, Y. S. Hwang, Y. H. Park, T. Ono and J. Y. Han (2018). “The dynamic development of germ cells during chicken embryogenesis.” Poult Sci 97(2): 650–657.

